# Proteomic characterization of neuronal extracellular vesicle interactomes in Alzheimer’s disease mouse model through TurboID-based proximity labeling

**DOI:** 10.1101/2025.10.22.684016

**Authors:** Brendan J. Gibbs, Yunfei Chen, Mohammad Abdullah, SonNgoc Nguyen, James McQuaker-Kajiwara, Bridgette Melvin, Alejandro Longtree-Preciado, Zhewei Liang, Maria Paz Gonzalez-Perez, Seiko Ikezu, Tsuneya Ikezu

## Abstract

Extracellular vesicles (EVs) are critical mediators of neuronal communication and have been implicated in propagating pathological processes in neurodegenerative diseases, including Alzheimer’s disease (AD). However, the molecular interactome of neuronal EVs *in vivo* remains poorly defined. Here, we employed TurboID-CD9–based proximity biotinylation to label and capture EV-interacting proteins in the hippocampus of wild-type (WT) and APP^NLGF^ knock-in AD mouse models. Adeno-associated viral delivery of hSyn1 promoter-driven TurboID-CD9 enabled neuron-specific EV tagging, followed by *in vivo* biotinylation and affinity purification of labeled proteins. Proteomic analysis using data independent acquisition liquid chromatography – mass spectrometry identified 5,502 proteins, with enriched pathways involving synaptic transmission, vesicle trafficking, and inhibitory neurotransmission. Comparative analyses revealed robust enrichment of GABAergic signaling components, including GABAA receptor subunits (Gabrb3, Gabra1, Gabbr2), Ncam1, and chloride transporters, in both WT and APP^NLGF^ EV interactomes, with additional disease-associated proteins (Mapt, Snca) and potassium channel enrichment observed in APP^NLGF^ mice. Proximity ligation assays validated direct EV-associated biotinylation of Ncam1, Gabrb3, and Gad1, with Gad1 showing significant upregulation in the APP^NLGF^ cohort. *In silico* HADDOCK docking supported stable interactions between CD9 and these target proteins, revealing plausible EV-protein interfaces. These findings define the *in vivo* neuronal EV interactome and its remodeling in amyloid pathology, implicating EV-associated GABAergic and ion channel proteins in network excitability regulation. This work establishes a proteomic and structural framework for understanding EV-mediated signaling in health and disease, providing candidate targets for therapeutic modulation of excitatory / inhibitory balance in AD.

## Introduction

A key regulatory and functional mechanism of all cells in the central nervous system (CNS) is the biogenesis, release, and integration of extracellular vesicles (EVs). EVs serve a diversity of functions in neuronal networks, from maintaining cellular homeostasis to serving as vehicles for the transfer of complex molecular signals ^1^. Although the function of EVs and multivesicular bodies (MVBs) in the lysosomal degradation pathways are well-established, emerging research underscores the significant impact of EVs on intercellular communication in the CNS ^2–4^. When EVs are released from a cell in the CNS, communication between cells can be established through the triggering of surface receptors leading to second messenger mediated signaling cascades or by delivering cell specific cargos consisting of proteins, nucleic acids and small molecules ^5,6^. For example in hippocampal neurons, heightened N-methyl-D-aspartate (NMDA) receptor activity leads to increased release of exosomes containing the transmembrane adaptor protein proline-rich 7 (PRR7). This activity-dependent EV release leads to neuronal uptake of exosomal PRR7 through membrane fusion which eliminates excitatory synapses in neighboring neurons by blocking the exosomal secretion of Wnts, activation of glycogen synthase kinase 3β, and promoting proteasomal degradation of postsynaptic density proteins ^7^. EV mediated interactions such as these highlight the dynamic influence of EVs on target cell function, however the role of EVs in modulating neuronal excitability and protein expression in the CNS is still not fully understood.

Emerging research also implicates EVs in the spread of pathological processes within the CNS. EVs released from neurons ^8^, astrocytes ^9^, microglia ^10^, and brain endothelial cells ^11^ have been shown to carry pathogenic proteins and small molecules across different neurodegenerative pathologies, ultimately causing mitochondrial dysfunction, oxidative stress, and inflammatory responses in recipient cells, accelerating neuronal injury and neurodegenerative cascades ^11^. In Alzheimer’s disease (AD), exosomes released from activated microglia help propagate neuronal tau pathology from the entorhinal cortex to the dentate gyrus exacerbating the spread of the disease ^12^. Additionally, exosomes secreted by microglia and neurons in Parkinson’s disease has been implicated in the spread of α-Syn leading to the neuronal death hallmarked by this pathology ^13,14^. Despite the importance of EVs in both homeostatic and pathological contexts, our knowledge of the targets and interactions of EVs and their cargo in the brain remains limited

For any comprehensive understanding of how EVs and their associated cargo interact with target cells regardless of homeostatic or pathological conditions, it’s essential to pinpoint the subcellular localization of EVs and their cargo in target cells. For this reason, imaging-based approaches, including fluorescent-based uptake studies, have been employed to elucidate the *in vivo* dynamics of EVs ^15,16^. Although these types of studies have been critical in furthering our understanding of how different CNS cells release and incorporate EVs ^17,18^, they generally offer little insight into the specific protein-protein interactions of EVs and their cargo. Additionally, while other approaches such as traditional proteomics and RNA transcriptomics have provided significant insights into the protein and RNA landscapes within individual cells ^19–21^, these methods often fail to pinpoint the specific origins or offer detailed spatial resolution of the proteins detected. Recent proteomic advancements however have enabled the exploration of protein-protein interactions through proximity-dependent labeling. Techniques like TurboID and APEX2 employ an engineered biotin ligase or peroxidase enzyme to label proteins within a 10 nm radius under physiological conditions ^22,23^, facilitating the study of specific protein interactomes. Although these groundbreaking techniques have been used to uncover key cell-type specific proteomic profiles in the CNS during different pathological states ^24^, these techniques have not been readily applied to understanding the dynamics of EV interactions in the brain. Leveraging these innovative methods, we aimed to elucidate the interactome of EVs from neurons and their associated cargo within the CNS of healthy and AD mouse models.

Employing the TurboID proximity labeling method, we aimed to decipher the interactions between neuron-derived EVs and CNS target cells under homeostatic and AD pathological states. We crafted an adeno-associated virus (AAV) to express hSyn1-mCherry-TurboID-CD9, enabling neuron-specific EV tagging. The AAV was administered intracranially into the dentate gyrus of both wildtype mice, for profiling the brain EV interactome in a healthy brain environment, and amyloid precursor protein (APP^NLGF^) mouse model to explore the EV interactome associated with AD.

## Methods

### Mouse models

All experiments were conducted in accordance with the Mayo Clinic Institutional Animal Care and Use Committee (IACUC) guidelines. C57BL/6 mice were purchased from Jackson Laboratory, and APP^NLGF^ mice were provided by Drs. T. Saito and T. Saido at RIKEN ^25^. All mice (WT C57BL/6 and APP^NLGF^) were housed in the vivarium managed by Mayo Clinic Animal Research with a 12 h light–dark cycle and ad libitum access to food and water. This study utilized 33 total mice, including 15 male C57BL/6 mice, 15 male APP^NLGF^ mice, and 3 female APP^NLGF^ mice, all 2 months old at the start of the experiment. Animals were randomly assigned to one of the following three experimental groups: TurboID + Biotin (TB), Saline + Biotin (SB), TurboID + Saline (T). Mice were bilaterally injected with AAV-hSyn1-mCherry-TurboID-CD9 or saline controls into the hippocampus, followed by intraperitoneal (IP) injections of biotin or saline starting on day 18 post-surgery.

### Generation of AAV-hSyn1-mCherry-TurboID-CD9 vector

TurboID-CD9 insert (1798 bp) and AAV-SYN1-mCherry vector backbone (4568 bp) were digested with BamHI and NotI, followed by gel purification using a Qiagen gel extraction kit after separation on 0.8% agarose gels. Purified DNA fragments were quantified using Nanodrop. The insert and vector were ligated at a 1:1 ratio using T4 DNA ligase, and 2 µL of the ligation mixture was transformed into 20 µL Sure-2 competent cells. After a 30-minute incubation on ice, cells were recovered in 200 µL SOC medium with shaking at 37°C for 1 hour and plated on LB agar containing 50 µg/mL ampicillin. Plates were incubated overnight (≤18 hours) at 37°C inverted. Individual colonies were inoculated into 4 mL LB with 50 µg/mL ampicillin and grown at 37°C with agitation for 12–18 hours. Positive clones were confirmed by BamHI/NotI digestion of miniprep DNA, verifying the presence of the 1798 bp insert and 4568 bp vector. Selected clones underwent endotoxin-free maxiprep with 500 mL LB culture to yield at least 200 µg plasmid DNA for AAV preparation. Expression of the target proteins was validated in N2a cells. Plasmids were then submitted for AAV production with the PHP.eB serotype at Boston Children’s Hospital.

### In vivo stereotaxic microinjections

AAV-hSyn1-mCherry-TurboID-CD9 or saline was bilaterally microinjected into the hippocampus of 8-week-old mice under general anesthesia. Anesthesia was induced with 5% isoflurane and maintained at 3% (v/v) using a continuous isoflurane delivery system. Anesthetic depth was monitored throughout the procedure. Mice were positioned in a stereotaxic frame (David Kopf Instruments) with their noses placed in a veterinary-grade anesthesia ventilation system (VetEquip), and their heads were stabilized using blunt ear bars. Pre-operatively, animals received subcutaneous injections of meloxicam (1 mg/kg) and 0.5 mL saline. Hair at the surgical site was trimmed with clippers, and the area was disinfected three times with alternating applications of 10% povidone iodine and 70% ethanol (v/v). A midline skin incision was made, and craniotomies (1–2 mm in diameter) were created bilaterally above the parietal cortex (-2.18 mm AP, ±1.55 mm ML) using a high-speed drill equipped with a small steel burr. Sterile saline was applied to the skull during drilling to minimize heat generation. Using a stereotaxic apparatus for guidance, bilateral injections were performed with a 10 μL Hamilton syringe (#701) at the following coordinates: -2.18 mm anterior to bregma, ±1.55 mm lateral to the midline, and 2.6 mm ventral to the pial surface. A syringe pump (SmartTouch Pump, World Precision Instruments) was used to deliver 1 μL of AAV, adjusted to a titer of 1.0 × 10¹³ genomcopies per mL in sterile 0.1 M PBS, at a controlled rate. After injection, the needle was left in place for 5 minutes before being slowly withdrawn to minimize reflux. Surgical wounds were closed using tissue adhesive (Vetbond, 3M). Post-operatively, animals were placed on a low-voltage heating pad until fully recovered from anesthesia. They were then transferred to clean cages containing DietGel and Hydrogel. Meloxicam (1 mg/kg) was administered subcutaneously once daily for 48 hours following surgery.

### In vivo TurboID protein biotinylation

Three weeks after AAV microinjection, mice were intraperitoneally injected once daily with 500 mg/kg biotin (Sigma-Aldrich, B4501) dissolved in sterile phosphate-buffered saline (PBS) for three consecutive days. The mice were sacrificed, and brain tissues were harvested 24 hours after the final biotin injection.

### Mouse Brain Tissue Collection and Preparation

For tissue intended for immunofluorescence (IF) staining, mice were anesthetized with isoflurane and transcardially perfused with 30 mL of PBS, followed by 20 mL of 4% paraformaldehyde (PFA). Brains were carefully extracted and post-fixed in 4% PFA overnight at 4°C. After fixation, the brains were cryoprotected in 30% sucrose until fully infiltrated. Tissues were then embedded in OCT compound, frozen, and sectioned for IF analysis. For tissue intended for biotinylated protein pull-down, mice were transcardially perfused with 30 mL of PBS, and the brains were immediately extracted and placed in ice-cold PBS. Cortical tissue anterior and posterior of the AAV injection site were isolated and weighed (approximately 100 mg per sample). The tissue was then transferred into 1 mL of RIPA lysis buffer (supplemented with Protease and Phosphatase Inhibitor Cocktail, Thermo Scientific, 78442) and finely minced on ice. Homogenization was performed using a dounce homogenizer to ensure efficient lysis. The homogenized tissue was subjected to sequential centrifugation steps to clarify the lysate: 500 × *g* for 10 minutes to remove large debris. 2,000 × *g* for 10 minutes to further clarify the supernatant.5,000 × *g* for 20 minutes, after which the final supernatant was carefully collected and used for biotinylated protein pull-down assays.

### Immunofluorescence Co-Staining

Mouse brain sections were subjected to immunofluorescence staining to visualize HA, mCherry, and biotinylated proteins using streptavidin. Brain tissues were fixed in 4% paraformaldehyde, cryoprotected in 30% sucrose, embed in oct, and sectioned at 30-μm thickness. Sections were permeabilized with 0.1% Triton X-100 in PBS, blocked with 5% normal serum, and incubated overnight at 4°C with primary antibodies against HA (3724S, Cell Signaling Technologies). Following primary antibody incubation, sections were washed and incubated with Alexa Fluor 488-conjugated secondary antibodies, along with Alexa Fluor 647-conjugated streptavidin (016-600-084, Jackson Immunoresearch) to detect biotinylated proteins. Sections were mounted with Fluoromount-G with DAPI. Images were captured using a laser-scanning confocal microscope (Leica SP8 with lightning) to confirm co-localization of HA, mCherry, and streptavidin signals.

### TurboID biotinylated protein pull-down

Biotinylated molecules were purified using Pierce Streptavidin Magnetic Beads (Thermo Scientific, 88816) following the manufacturer’s instructions. Briefly, 50 μL of Pierce Streptavidin Magnetic Beads were thoroughly mixed and pre-washed three times with 1 mL of binding/wash buffer (Tris-buffered saline, TBS, pH 7.5, with 0.1% Tween-20) using a magnetic stand. The final supernatant from brain tissue lysates was combined with 10 μg of biotin monoclonal antibody (Rockland, 200-301-098) and incubated overnight at 4°C with gentle mixing. The antibody-antigen mixture was then added to the pre-washed magnetic beads and incubated for 2 hours at room temperature with gentle mixing. The beads were washed three times with binding/wash buffer to remove non-specific binding. Bound proteins were eluted by incubating the beads with 100 μL of SDS-PAGE reducing sample buffer at 96–100°C for 5 minutes, followed by magnetic separation. The eluted supernatant, containing the target antigen, was collected for LC-MS analysis. Protein concentrations of the samples were determined using the Micro-BCA Protein Assay Kit (Thermo Scientific, 23235).

### Western Blotting for Detection of Biotinylated Molecules

Homogenate samples (final supernatant from brain tissue lysates) were mixed with SDS-PAGE reducing sample buffer and heated at 96–100°C for 5 minutes to denature the proteins. A total of 4 μg of homogenate sample and 1 μg of purified biotinylated molecules were loaded onto a 4– 20% polyacrylamide precast gel (Bio-Rad, 4561096) for protein separation. Proteins were transferred onto a nitrocellulose membrane (0.45 μm pore size, Bio-Rad) using a wet transfer system at 250 mA for 1 hour on ice. Membranes were blocked with 5% non-fat milk in Tris-buffered saline containing 0.1% Tween-20 (TBST) for 1 hour at room temperature to prevent non-specific binding. The membranes were then incubated with streptavidin-HRP conjugate (Thermo Scientific, 21140) diluted in 3% BSA in TBST for 1 hour at room temperature with gentle agitation. After incubation, the membranes were washed three times with TBST for 10 minutes each. Biotinylated molecules were visualized using enhanced chemiluminescence (ECL) reagent (EMD Millipore, WBKLS0500) and detected using the Bio-Rad ChemiDoc Imaging System.

### LC-MS/MS

The samples (4 µL injection) were analyzed on a TimsTOF Pro2 (Bruker) mass spectrometer, which was coupled to a nanoElute LC system from Bruker. Peptides were then loaded and separated on an in-house-made 75 μm I.D. fused silica analytical column packed with 25-cm ReproSil-Pur C18-AQ (Dr. Maisch, GmbH, 120 Å, 3 μm) particles to a gravity-pulled tip. A 60-minute and 30-minute gradient was employed for DDA-PASEF and DIA-PASEF, respectively, with a flow rate set at 500 nL/min. The captive nano-electrospray voltage was maintained at 1600V, using one column configuration (no trap) and a solvent composition of Solvent A: 0.1% FA in water and Solvent B: 0.1% FA in ACN.

DDA-PASEF method: The method consists of 10 MS/MS PASEF scans per topN acquisition cycle, with ramp and accumulation times of 100 ms, covering an m/z range from 100 to 1700 and an ion mobility range (1/K0) from 0.70 to 1.30 V s/cm^2^. Collision energy settings followed a linear function of ion mobility, ranging from 20 eV at 0.6 V s/cm^2^ to 59 eV at 1.6 V s/cm^2^, using default parameters. Calibration of the instrument was performed using three ions from the ESI-L Tuning Mix (Agilent) (m/z 622, 922, 1222).

Dia-PASEF method: Samples were acquired with a method consisting of 22 mass width windows (40 Da width, from 275 to 1155 Da) with 1 mobility windows each covering the ion mobility range (1/K_0_) from 0.70 to 1.30 V s/cm^2^. These windows were optimized with the Window Editor utility from the instrument control software (timsControl, Bruker) using one DDA-PASEF run acquired of the analyzed samples. Collision energy setting and calibration were the same as DDA-PASEF analysis.

### LC-MS/MS Data Analysis Workflows

Fragpipe/MSFragger/ScaffoldQ+S DDA label free quantitative analysis: The custom workflow ‘LFQ-MBR’ from FragPipe version 21.1 was used, including database search with MSFragger (version 4.0), deep-learning prediction rescoring with MSBooster, Percolator and ProteinProphet (Philosopher version 5.1.0) for PSM validation and protein inference. The raw .d files from Bruker were searched against a isoform fasta database (UP000000589) appended with common contaminant proteins. Decoy reversed sequences were appended to the search database. The default MSFragger search parameters were used, except precursor and fragment mass tolerances were set to 20 ppm; enzyme: trypsin; peptide length 7-35. Variable modifications were set to oxidation of Met and acetylation of protein N-t, and fixed modifications to carbamidomethylation of Cys. Percolator and ProteinProphet default options were used, and results were filtered by 1% FDR at protein level. The searched results were then imported to Scaffold Q/S 5.2.2 for data visualization and statistical analysis.

Spectronaut, Library-free DIA analysis (directDIA+): The directDIA+ workflow in Spectronaut (v18.2) was used for analyzing the diaPASEF dataset with no need to build a library previously from the DDA runs. The raw .d files from Bruker were converted to HTRMS files automatically right after acquisition via Spectronaut HTRMS Converter program. Data were searched against a reviewed mouse isoform fasta database (UP000000589). The default factory settings were used for the Pulsar search and DIA analysis (including Trypsin/P as enzyme, 7-52 peptide length range; Oxidation of Me and acetylation of Protein N-t as variable modifications, carbamidomethyl of C as fixed modification, and 1% FDR for PSM, peptide and protein identification). Automatic cross run normalization and Protein LFQ method were used for protein quantification, and quantity was determined at MS2 level using the area of extracted chromatogram traces.

### Bioinformatics Pipeline

Data-Independent Acquisition (DIA) is a mass spectrometry approach in which all peptides detected at the MS1 level can be fragmented and analyzed at the MS2 level. Differential expression (DE) analysis in DIA proteomics aims to identify proteins whose abundances differ significantly between control and case groups.

Using Spectronaut to process the DIA data, a total of 5,502 proteins were identified. To filter out mitochondrial proteins, we applied keyword filtering (e.g., “mitochondrial,” “dehydrogenase,” “carboxylase”), which resulted in a refined dataset of 5,098 proteins.

For sample organization, all groups each contained three technical replicates and each biological group comprised six samples. To streamline the analysis, replicate measurements were averaged, yielding one value per sample. Pairwise DE comparisons were performed across the groups (WT-TB vs. WT-SB, APP-TB vs. APP-SB, and APP-TB vs. WT-TB) using the in-browser DIA-Analyst tool. Proteins were considered differentially expressed if the absolute log2 fold change (|log2FC|) was ≥ 1 and FDR < 0.05. Missing values were imputed using a Perseus-type approach ^26^.

### Enrichment analysis

Enrichment analysis was performed using custom Python scripts (utilizing packages such as Matplotlib, Pandas, and NumPy). A Z-score was calculated for each protein to assess differential abundance, and proteins were ranked based on a combined significance score incorporating both log2 fold change (log2FC) and statistical significance (−log10 *p*-value). Differentially enriched proteins (p < 0.05, log2(FC) > 1) underwent GO Biological Pathways analysis using Enrichr ^27^ and STRING analysis ^28^ was used to understand protein-protein interactions of enriched pathways. Cell-specific analysis of differentially enriched proteins was done by the comparison of enriched protein datasets with GSEA (MSigDB) ^29^ and PanglaoDB ^30^ databases.

### Proximity Ligation Assay

Proximity ligation assays (PLA) were used to validate our proteomic findings by investigating protein–protein interactions. Brightfield PLA was performed according to the manufacturer’s protocol (#DUO92001 and #DUO92005, Sigma–Aldrich). Brain sections from our TB, SB and T groups in both WT and APP^NLGF^ mice were fixed with 4% paraformaldehyde and washed with PBS. The sections were blocked with Duolink Blocking Solution for 1 hour at 37 °C, and the slices were subsequently incubated with anti-rabbit streptavidin (0437R, Bioss Antibodies) and anti-mouse target protein Gad1 (MAB5406B, Sigma-Aldrich), Gabrb3 (ab98968, Abcam), Ncam1 (ab9018, Abcam)) primary antibodies overnight in a humid chamber at 4 °C. The following day, the slices were incubated with the secondary PLA probes anti-mouse PLUS and anti-rabbit MINUS in a preheated humid chamber for 1 hour at 37 °C. After washing, the sections were ligated (#DUO92007, Sigma–Aldrich) for 30 minutes and amplified with polymerase for 100 minutes at 37 °C. The sections were visualized by the addition of Duolink In Situ Mounting Medium with DAPI and imaged using a confocal microscope under consistent laser and gain settings. Signal analysis was performed in cortical regions overlying the hippocampus and manually quantified using IMARIS 3D rendering.

### In-silico Docking Simulation

Three-dimensional (3D) structural data and structure prediction of biomolecular interactions for proteins were generated using AlphaFold 3 ^31^. Predicated Template Modeling (scores ≥ 0.5) was assessed alongside predicted Local Distance Difference Test (scores ≥ 70) to filter model quality to ensure models had well-defined domain boundaries for downstream analyses. Subsequently, models were superimposed upon experimentally published structures on UniProt, with preference given to the highest-resolution Cryo-EM, NMR, or X-ray crystallographic entries available. Structural alignments were performed in ChimeraX 1.10.1. ^32^ using the MatchMaker function and pruned root-mean-square deviation (RMSD) values were calculated. Models exhibiting RMSD ≤ 1.1 Å relative to their experimental counterparts, advanced to docking using HADDOCK 2.4. ^33^. Docking was performed ab initio using HADDOCK 2.4. without specifying active or passive residues. The number of runs generated for it0, it1, and itw were set to 10,000, 600, 600, respectively. Centre-of-mass restraints were applied, and clustering with a minimum cluster size of 2. Runs were ranked based on the weighted HADDOCK composite score; cluster size, RMSD from the overall lowest-energy structure, Van der Waals energy, electrostatic energy, desolvation energy, restraints violation energy, buried surface area, Z-score.

Visualization and Polar Contact Mapping Docking geometries were visualized in PyMOL ver. 3.1.6.1. (Schrödinger, LLC) To identify interfacial polar contacts, residues within 3 Å were selected using logical “within” operators. Polar contacts were visualized as dashed yellow lines via the “distance” command (dist polar_contacts, CD9, GABRB3, cutoff=3). The sequence viewer in PyMOL 3.1.6.1. was used to identify and annotate residues involved in these contacts. Complementary visualization was performed in ChimeraX 1.10.1. for figure creation of the docking interface and highlight contact sights from PyMOL 3.1.6.1.

## Results

### Neuronal CD9 interactome in the brain of healthy WT mice

We set out to comprehensively classify the interactome of neuronal EVs in the WT murine brain to better understand neuronal EV secretion and integration *in vivo* (Fig. 1A). We first analyzed the proteomic profiles of all three of our WT groups to understand similarities and differences in protein expression between groups. We observed that our WT TurboID + Biotin (WT-TB) group had significant Principal Component Analysis (PCA) separation (Fig. 1B), highest number of captured proteins (Fig. 1C) and the highest number of unique proteins (Fig. 1D) compared to our two control groups: TurboID + Saline (WT-T) and Saline + Biotin (WT-SB). We also observed 3559 shared genes across all groups, likely representing endogenously biotinylated proteins in the healthy murine brain. We next compared protein expression profiles by log2-centered heatmap analysis of protein intensities between all groups (Fig. 1E), which revealed distinct expression patterns in the TB condition, helping further support the successful targeting and labeling of neuronal EV-associated proteins through our hSyn1-CD9 TurboID vector. To investigate the proteins involved in the neuronal EV interactome under biotin-supplemented conditions, reflecting neuronal EV synaptic dynamics, we compared expression profiles of the WT-TB group against the WT-SB control condition. We first ranked proteins based on combined significance, incorporating both log2 fold change (log2FC) and statistical significance (−log10 *p*-value). Z-scores were then calculated to compare protein abundances between the TB and SB datasets, allowing us to identify proteins that were significantly enriched in the TB condition by quantifying how strongly individual protein levels deviated from the background distribution. Ranked proteins were then plotted against their respective Z-score, allowing us to identify the top enriched proteins in the WT-TB condition (Fig. 1F). Through this approach, we identified several proteins implicated in essential EV-related and neuronal processes, including Mapt, microtubule-associated protein; Septin11, a cytoskeletal regulator involved in vesicle trafficking; Ncam1, a key mediator of neuronal cell-cell interactions and synaptic plasticity; Gabrb3, a subunit of GABA_A_ receptor complexes involved in inhibitory neurotransmission and Cd47, a membrane protein linked to immune modulation and synaptic pruning. Next, to better characterize significantly enriched proteins in the TB dataset, we performed a volcano plot analysis, comparing fold change to statistical significance to identify proteins with both high differential abundance and statistical significance between the WT TB dataset and the SB control (Fig. 1G). Enriched proteins identified from this analysis were then compared against volcano analysis between our control groups (WT T vs. SB) (Supplementary Fig. 2B–C) to isolate proteins uniquely enriched in the TB condition. Among the uniquely enriched proteins in the TB dataset, we observed prominent upregulation of components involved in vesicle trafficking and endocytosis, including Sirpa, a membrane receptor that modulates phagocytic signaling and Lingo1, a transmembrane protein known to regulate endocytosis, axonal growth, and synaptic remodeling. We also observed enrichment of proteins involved in synaptic transmission and neuronal excitability including Gabrb3, a subunit of the GABA_A_ receptor complex central to inhibitory neurotransmission, and Erbin, a postsynaptic scaffolding protein implicated in stabilizing receptor complexes and regulating synaptic architecture. Additionally, key neuronal transporters such as Slc6a6, a taurine transporter involved in neuromodulation and osmotic balance, were observed. Lastly, several proteins linked to G protein–coupled receptor (GPCR) signaling were enriched in the TB dataset. This included Gpr158, an orphan GPCR implicated in synaptic organization, and Shb, an adaptor protein that modulates downstream signaling cascades downstream of receptor tyrosine kinases and GPCRs. We further classified the most enriched proteins in our WT-TB experimental group through GO biological pathways analysis and found a diverse range of proteins across various cellular functions and pathways (Fig. 1H; Supplementary Table). We observed the significant enrichment in pathways such as the gamma-aminobutyric acid (GABA) signaling pathway (GO:0007214; e.g. Gabbr2, Gabra1), chemical synaptic transmission (GO:0007268; e.g. Gprin1, Amph), and inorganic cation transmembrane transport (GO:0098662; e.g. Slc30a1, Slc8a1).

**Figure 1:**
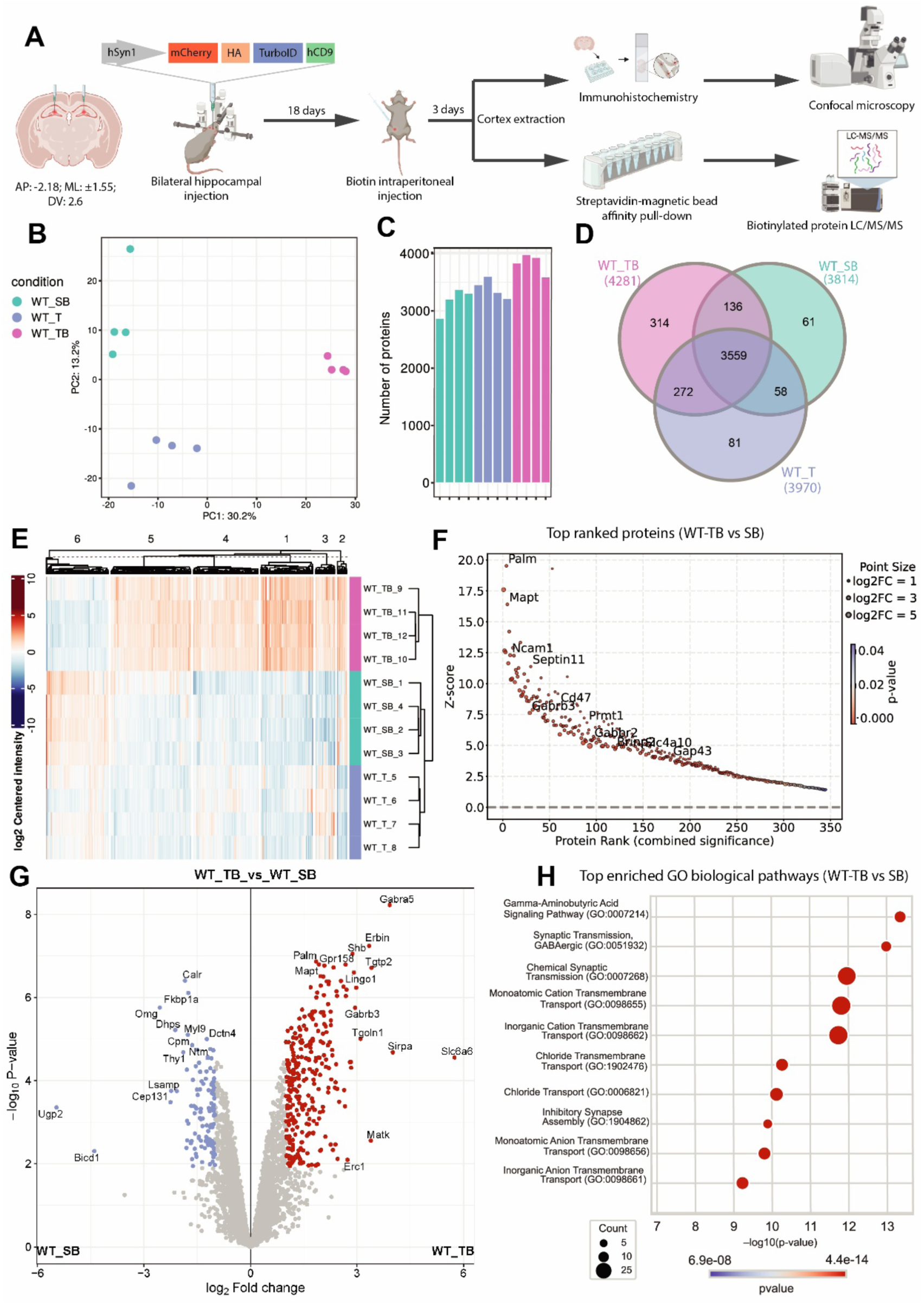
Comprehensive proteomic analysis of CD9 interactome in WT mice. (**A**) Schematic overview of TurboID proteomic workflow. **(B)** Principal component analysis (PCA) of the TB experimental group against the SB (top) and T (bottom) control groups. **(C)** Number of proteins detected in each of our three groups. **(D)** Venn diagram showing protein overlaps across groups. **(E)** Heat map of protein expression across groups. (**F**) Top ranked proteins (combined significance) of the TB group when compared against the SB control plotted against Z-score. (**G**) Volcano plot showing enriched proteins in the TB group compared to controls. Comparison includes WT-TB vs. WT-SB, with significance thresholds set at one-way ANOVA, FDR < 0.05, log2 FC > 1, and log2 FC < -1. (**D**) GO biological pathway analysis of enriched proteins in TB vs SB groups.

### Neuronal CD9 interactome in the brain of an Alzheimer’s mouse model (APP^NLGF^)

We next set out to comprehensively classify the interactome of neuronal EVs in APP^NLGF^ mice to understand EV interactions in amyloid pathology. We first analyzed the proteomic profiles of our APP^NLGF^ groups to determine shared and unique proteins between conditions and found that the APP^NLGF^ TurboID + Biotin (APP-TB) group had significant PCA separation (Fig. 2A), highest number of captured proteins (Fig. 2B) and the highest number of unique proteins (Fig. 2C) compared to our two control groups: TurboID + Saline (APP-T) and Saline + Biotin (APP-SB). We observed 4982 shared genes across all groups, likely representing endogenously biotinylated proteins in the amyloid pathological brain. Next, we compared protein expression profiles of all groups by log2-centered heatmap analysis of protein intensities (Fig. 2D), which revealed distinct expression patterns in the TB condition, helping further support the success of our TurboID vector in APP^NLGF^ mice. To investigate the proteins involved in the neuronal EV interactome during amyloid pathology, we compared expression profiles of the APP-TB group against the APP-SB control condition. We first ranked proteins based on combined significance, incorporating both log2 fold change (log2FC) and statistical significance (−log10 *p*-value). Z-scores were then calculated to compare protein abundances between the TB and SB datasets. Ranked proteins were then plotted against their respective Z-score, allowing us to identify the top enriched proteins in the APP-TB condition (Fig. 2E). Through this approach, we identified proteins implicated in essential EV-related and synaptic processes, including Bin1, involved in membrane remodeling and endocytosis; Snca, involved in synaptic function and linked to protein aggregation in AD; Mapt, tau protein heavily implicated in AD; Ncam1, a key mediator of neuronal cell-cell interactions and synaptic plasticity; Gabrb3, a subunit of GABA_A_ receptor complexes involved in inhibitory neurotransmission; and Plxna4, a receptor involved in axonal processes and neural circuit formation. To better understand the neuronal EV interactome in AD pathology, we further characterize proteins significantly enriched in the TB dataset through volcano plot analysis comparing fold change with statistical significance to identify proteins exhibiting both substantial differential abundance and significance between the APP TB dataset and the SB control (Fig. 1G). Enriched proteins identified from this analysis were then compared against volcano analysis between our control groups (APP T vs. SB) (Supplementary Fig. 2D–E) to highlight proteins uniquely enriched in the TB condition. Among the uniquely enriched proteins in the TB condition, we observed increased enrichment of proteins involved in neurotransmission and excitability including Gabrb3, a GABA_A_ receptor subunit involved in inhibitory neurotransmission; Kcnma1, a large-conductance calcium-activated potassium channel regulating neuronal excitability; Kcnk2, a two-pore-domain potassium channel modulating resting membrane potential; Dpp6, which modulates A-type potassium channel kinetics; and Cplx2, a synaptic protein involved in neurotransmitter release. Additionally, neuronal adhesion molecules Ncam1 and Ncam2, implicated in synaptic plasticity and endocytosis, were upregulated, and Cd47, associated with vesicle trafficking and immune signaling. Lastly, neurodegeneration-related proteins such as Mapt and Sncb were also enriched. Using GO biological pathway analysis, we further classified the most enriched proteins in the APP-TB group (Fig. 2G; Supplementary Table) and identified significant enrichment in pathways including Gamma-Aminobutyric Acid Signaling Pathway (GO:0007214; e.g. Gabra2; Gabrb3), Chloride Transmembrane Transport (GO:1902476; e.g. Slc6a1; Slc12a6) and Synaptic Vesicle Endocytosis (GO:0048488; e.g. Stx1a; Vamp2).

**Figure 2:**
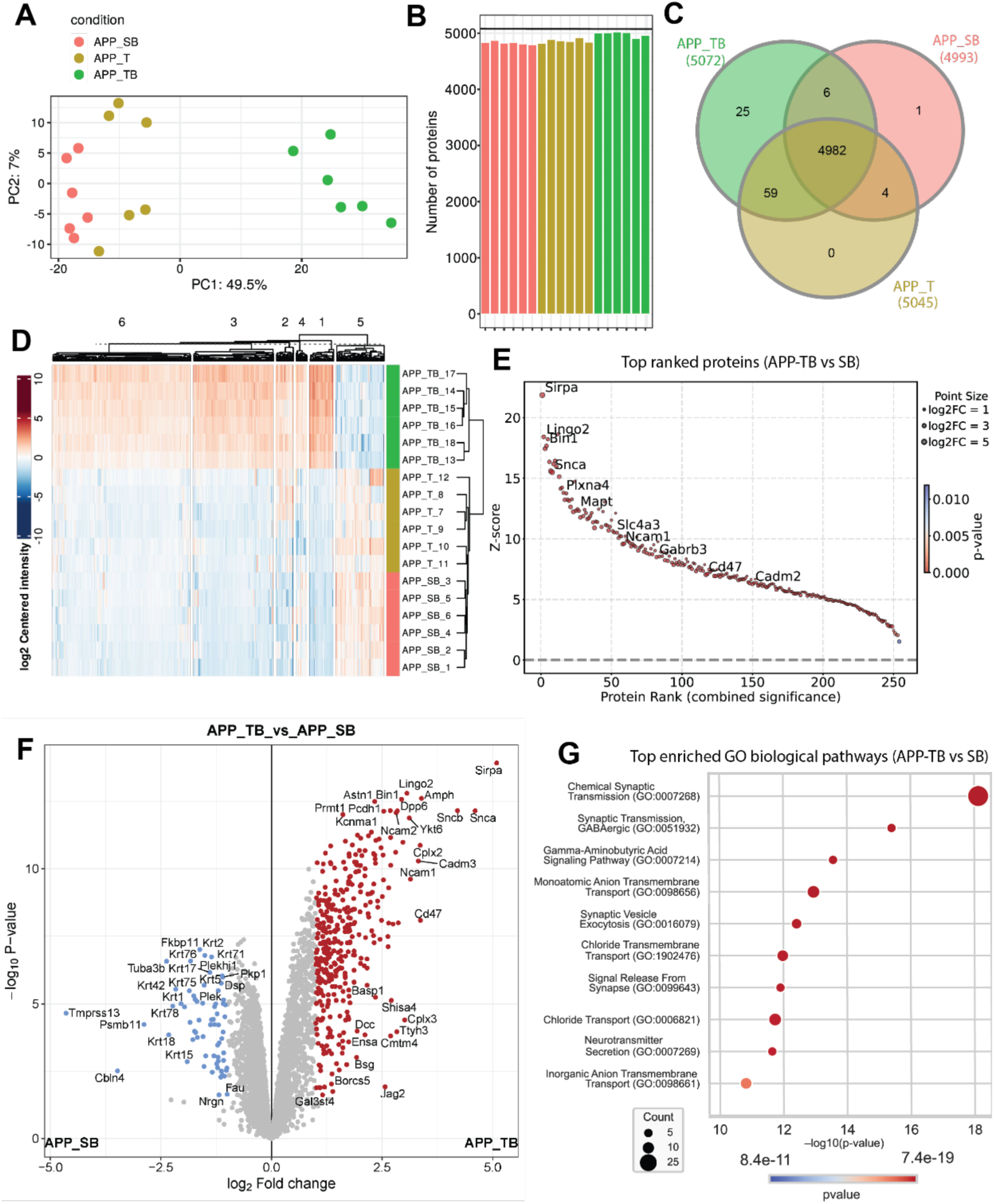
Comprehensive proteomic analysis of CD9 interactome in APP^NLGF^ mice. (**A**) PCA of the TB experimental group against the SB (top) and T (bottom) control groups. **(B)** Number of proteins detected in each of our three groups. **(C)** Venn diagram showing protein overlaps across groups. **(D)** Heat map of protein expression groups. **(E)** Top ranked proteins of the TB group when compared against the SB control plotted against Z-score. (**F**) Volcano plot showing enriched proteins in the TB group compared to controls. Comparison includes APP-TB vs. APP-SB, with significance thresholds set at one-way ANOVA, FDR < 0.05, log2 FC > 1, and log2 FC < -1. (**G**) GO biological pathway analysis of enriched proteins in TB vs SB groups.

### Comparison of the CD9 interactome in WT and APP^NLGF^ mouse models

To understand how the EV interactome is altered in the context of amyloid pathology compared to homeostatic conditions, we compared the APP^NLGF^ and WT groups. We first directly compared enriched protein datasets between the APP-TB and WT-TB groups, finding 155 proteins shared, with 190 proteins uniquely enriched in the WT-TB group and 99 unique to the APP-TB condition (Fig. 3A). Additionally, PCA analysis revealed significant separation between the APP and WT TB conditions (Fig. 3A). Next, to avoid batch effect bias caused by differences in the total number of detected proteins between the WT and APP groups, we compared enriched pathways rather than direct proteomic comparison to allow us to better understand how the EV interactome may be altered in AD pathology. We first compared enriched (TB vs. SB) GO biological pathways between the WT and APP TB conditions to identify the top shared pathways, aiming to uncover robust proteomic interactions of neuronal EVs (Fig. 3B; Supplementary Table). Using GO biological pathway analysis, we identified top shared pathways between WT and APP TB conditions, including Chemical Synaptic Transmission (GO:0007268; e.g., Gria1, Gria2, Snap25), Gamma-Aminobutyric Acid Signaling Pathway (GO:0007214; e.g., Gabrb3, Gabbr2, Gabra1), Synaptic Vesicle Exocytosis (GO:0016079; e.g., Stx1b, Prrt2, Cplx2), and Chloride Transmembrane Transport (GO:1902476; e.g., Slc12a5, Slc6a1, Slc12a6). To gain deeper insight into the protein interactome within one of the top shared pathways, Gamma-Aminobutyric Acid Signaling Pathway (GO:0007214), we conducted STRING network analysis (Fig. 3C). This revealed a tightly connected network of enriched proteins, including GABA_A_ receptor subunits (Gabrb3, Gabbr2, Gabra1) and chloride transporters (Slc12a5, Slc6a1, Slc12a6), interacting closely with key regulatory proteins such as Hap1, involved in intracellular trafficking, and Gad1, the enzyme critical for GABA synthesis, highlighting the complexity of EV associated protein interactions in the context of inhibitory signaling. Next, we sought to characterize differences between the WT and APP TB conditions by comparing the enriched (TB vs SB) datasets to identify unique proteins in each condition. GO biological pathway analysis of uniquely enriched proteins in the WT TB condition included Axon Guidance (GO:0007411; Cntn4; L1cam), Chemical Synaptic Transmission (GO:0007268; Unc13c; Grin3a), Neuron Projection Morphogenesis (GO:0048812; Shtn1; Gpm6a) and Protein Phosphorylation (GO:0006468; Mink1; Matk) (Fig. 3D; Supplementary Table). To gain deeper insight into the protein interactome in the most enriched unique WT-TB pathway, STRING network analysis was done on proteins associated with the Axon Guidance pathway. This analysis revealed a coherent network of interactions between uniquely enriched proteins such as L1cam and Nrcam, both of which are critical cell adhesion molecules involved in axonal pathfinding and synapse formation. These molecules were shown to interact with Nrp1 and Sema3a, key guidance cue signaling components that mediate growth cone collapse and axonal repulsion (Fig. 3E). In the APP-TB condition, GO biological pathway analysis of unique enriched proteins revealed pathways including Action Potential (GO:0001508; Slc4a3; Kcnk2), Membrane Depolarization During Action Potential (GO:0086010; Hcn2; Scn2b), Potassium Ion Transport (GO:0006813; Slc24a4; Kcnq2), and Long-Term Synaptic Potentiation (GO:0060291; Gria3; Vamp2) (Fig. 3F; Supplementary Table). STRING analysis of the most enriched pathway, Action Potential, revealed interactions among uniquely enriched proteins directly involved in excitability and membrane potential dynamics such as Scn2b, Kcna4, and Kcnk2 with Sema8a, a semaphorin family member that can modulate voltage-gated ion channel activity and axon excitability (Fig. 3G). Lastly, we compared the enriched protein datasets (TB vs. SB) from WT and APP groups to single-cell marker databases to define the cell-type-specific interactome of neuron-derived EVs in each condition. This analysis revealed the presence of proteins associated with excitatory neurons, inhibitory neurons, astrocytes, microglia, oligodendrocytes, and Schwann cells in both WT (Fig. 3H) and APP (Fig. 3I) groups, underscoring the diversity of cellular targets by neuronal EVs and highlight the complexity of EV-mediated intercellular communication within the CNS.

**Figure 3:**
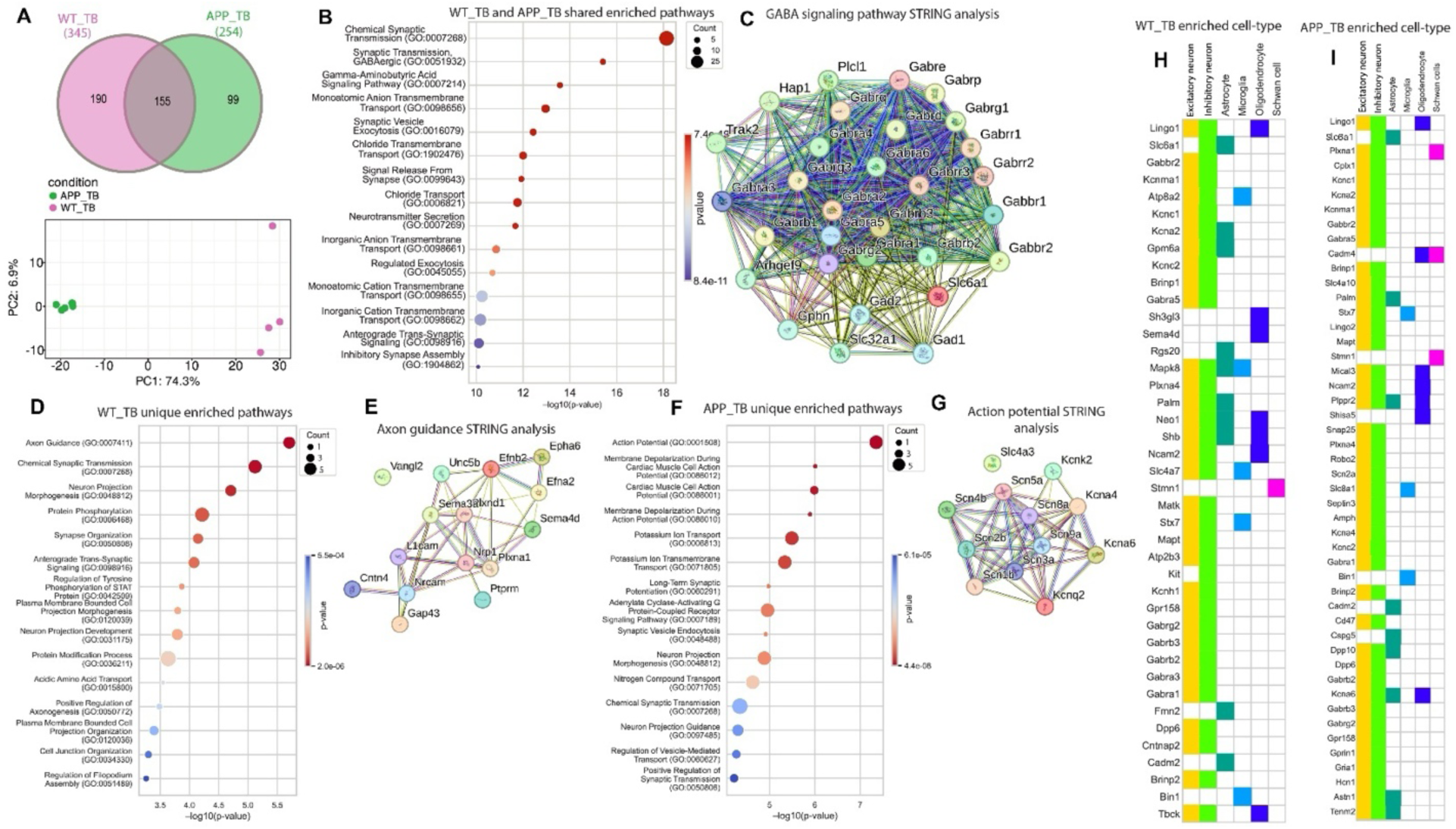
Comprehensive Proteomic Analysis: Comparison of CD9 Interactome in APP^NLGF^ Mice vs. WT Mice. (**A**) Venn diagram of enriched protein overlaps in APP-TB vs WT-TB mice. PCA comparison of APP-TB and WT-TB groups. (**B**) Top shared enriched GO biological pathways between WT-TB and APP-TB groups. **(C)** STRING analysis of enriched proteins in shared GABAergic synaptic transmission pathway. (**D**) Unique enriched GO biological pathways in the WT-TB group. **(E)** STRING analysis of enriched proteins in unique axon guidance pathway. (**F**) Unique enriched GO biological pathways in the APP-TB group. **(G)** STRING analysis of enriched proteins in unique regulation of action potential pathway. **(H)** Cell-type specific analysis of enriched proteins in the WT-TB group. **(I)** Cell-type specific analysis of enriched proteins in the APP-TB group.

### Proximity Ligation Assay (PLA) validation of target molecules

We next aimed to identify and validate key proteins of interest from our enrichment analysis using proximity ligation assays (PLA). Candidate proteins (TB vs. SB) were selected based on a combination of statistical significance, z-score ranking, and high abundance (imputed average >10) (Fig. 1F-G; Fig. 2E-F). These candidates were then compared to proteins enriched in our TurboID + Saline control dataset (T vs. SB) (Supplementary Fig. 2) to identify proteins uniquely enriched in WT and APP TB groups relative to controls (T and SB). The resulting enriched protein datasets from WT and APP TB conditions were cross-compared to identify top shared hits, including Mapt, Ncam1, and Gabrb3 (Fig. 1F-G; Fig. 2E-F). Among the top candidates, we prioritized the GABA_A_ receptor subunit Gabrb3 to investigate how EVs may modulate inhibitory neurotransmission, and Ncam1 to explore EV-mediated intercellular communication, including uptake and incorporation into recipient cells via endocytic pathways.

We also aimed to identify and validate a protein differentially expressed in the APP-TB condition relative to WT-TB that may be relevant to Alzheimer’s disease (AD) pathology. Given the shared enrichment of GABAergic signaling components, including nine GABA_A_ receptor subunits and the GABA transporter Slc6a1 (GAT1), between WT and APP conditions (Fig. 1H; Fig. 2G; Fig. 3B), we investigated whether any interacting partners of these shared proteins might be differentially expressed in APP-TB. STRING analysis of the shared GABAergic pathway proteins revealed strong interaction networks, notably with Gad1 (Fig. 3C), a key enzyme in GABA synthesis and a central regulator of excitatory/inhibitory (E:I) balance. We then compared Gad1 expression across WT-TB and APP-TB conditions relative to SB controls and observed a marked difference: while WT-TB vs. SB showed no significant change (p = 0.432; Log₂FC = –0.125), APP-TB vs. SB demonstrated significant upregulation (p = 0.042; Log₂FC = 0.289) (Supplementary Fig. 3D). Based on this finding, we further investigated whether neuronal EV–Gad1 interactions differ between WT and APP conditions.

To evaluate the biotinylation of these target proteins by TurboID, we performed PLA experiments for streptavidin in complex with Ncam1, Gabrb3, and Gad1 across WT and APP^NLGF^ groups to validate proteomic findings. For Streptavidin–Ncam1, APP-TB mice exhibited significantly elevated puncta relative to APP-SB (1793 vs 676; p = 0.001), while WT-TB also significantly differed from WT-SB (1257 vs 445; p = 0.031). No significance was observed between APP-TB and WT-TB groups (p = 0.27) (Fig. 4B, C). For Streptavidin–Gabrb3, both WT-TB and APP-TB groups showed significantly higher puncta than their respective SB controls (WT: 1500 vs 313; p = 0.02; APP: 1526 vs 421; p = 0.0003). No significance weas observed between APP-TB and WT-TB (p = 0.94) (Fig. 4E, F). For Streptavidin–Gad1, APP-TB mice showed a significant increase in PLA puncta compared to both APP-SB (2118 vs 397; p = 0.001) and WT-TB (2118 vs 499; p = 0.001), indicating enhanced labeling of GAD1 in the APP-TB condition. No significant difference was observed between WT-TB and WT-SB (499 vs 317; *p* = 0.31) (Fig. 4H, I).

**Figure 4:**
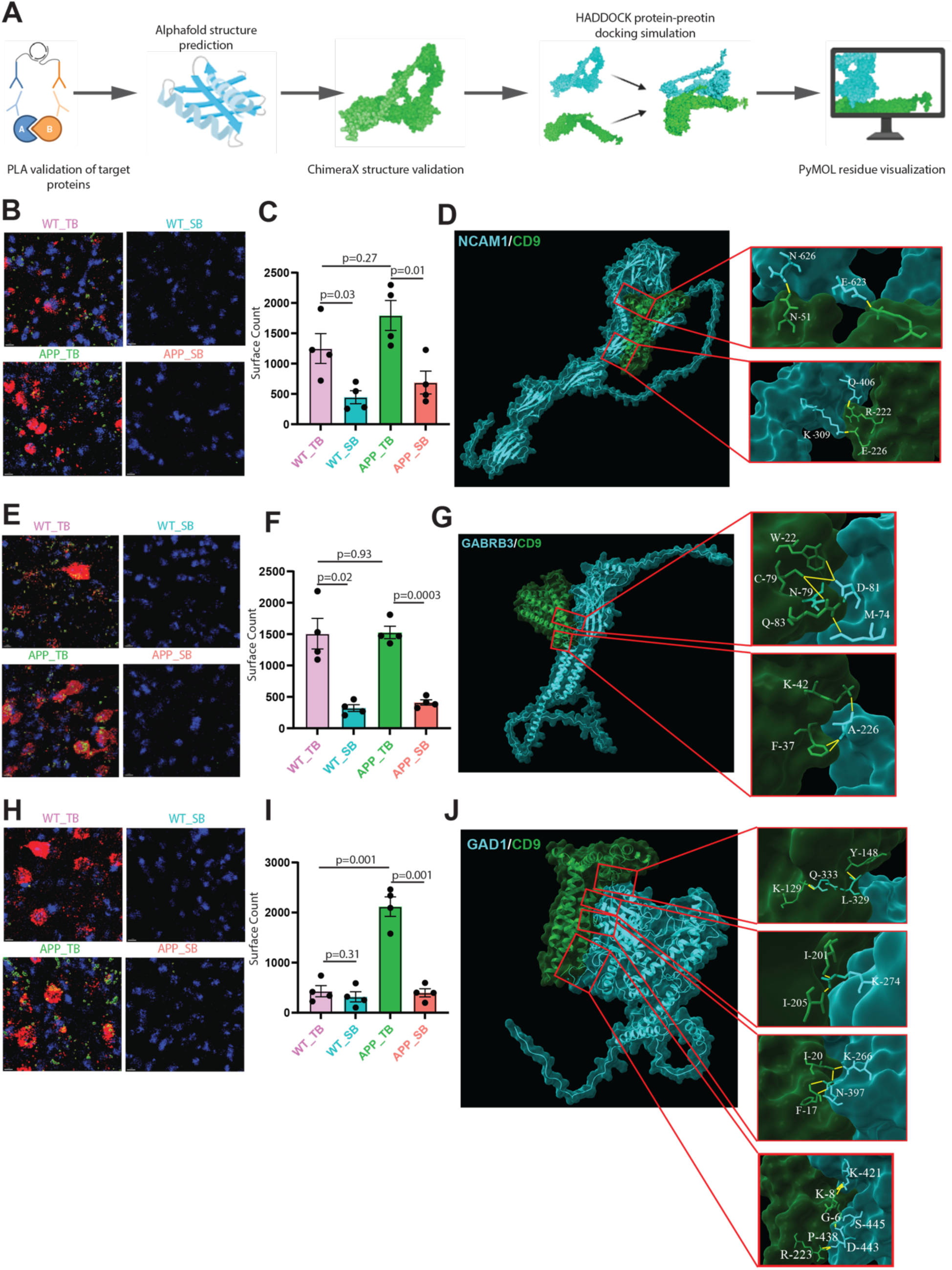
Proximity ligation assay (PLA) validation of biotinylation in proteins of interest. **(A)** Schematic overview of PLA validation and *in silico* modeling workflows of CD9 interactions with of proteins of interest. **(B)** PLA conjugation of Ncam1 with streptavidin in the TB and SB groups in both WT and APP^NLGF^ mice where mCherry represents TurboID infected neuron, blue represents DAPI and green represents PLA conjugation. **(C)** Comparison of PLA conjugation (surface count) of Ncam1 with streptavidin between all groups (One-way ANOVA). **(D)** HADDOCK modeling of CD9 protein-protein docking with Ncam1 where zoomed in images highlight specific residue interactions. **(E)** PLA conjugation of Gabrb3 with streptavidin in the TB and SB groups in both WT and APP^NLGF^ mice. **(F)** Comparison of PLA conjugation (surface count) of Gabrb3 with streptavidin between all groups (One-way ANOVA). **(G)** HADDOCK modeling of CD9 protein-protein docking with Gabrb3 where zoomed in images highlight specific residue interactions. **(H)** PLA conjugation of Gad1 with streptavidin in the TB and SB groups in both WT and APP^NLGF^ mice. **(I)** Comparison of PLA conjugation (surface count) of Gad1 with streptavidin between all groups (One-way ANOVA). **(J)** HADDOCK modeling of CD9 protein-protein docking with Gad1 where zoomed in images highlight specific residue interactions.

### In Silico Docking of CD9 with proteins of interest

To evaluate protein–protein interactions between neuronal EVs (CD9) and our validated proteins of interest, we performed HADDOCK docking simulations to gain insight into their potential binding interfaces and interaction stability (Fig. 4A). HADDOCK docking of CD9 with NCAM1 yielded a top cluster (Cluster 48) with a HADDOCK score of –90.5 ± 11.1, supporting a moderately stable interaction. The cluster had a size of 2, with an RMSD of 18.2 ± 0.1 Å, indicating consistent conformational sampling within the cluster. Key residues within 3 Å at the predicted interface included NCAM1: 266, 274 and CD9: 51, 68, 119, 191, 222, 226, suggesting a contact surface involving extracellular domains of both proteins (Fig. 4D). Energetically, the interaction was driven by favorable electrostatic energy (–160.7 ± 26.1 kcal/mol) and van der Waals contributions (–36.8 ± 4.1 kcal/mol), along with a buried surface area of 1727.0 ± 239.7 Å², consistent with a moderately extensive protein–protein interface. No restraint violations were observed, and the Z-score of 0.2 reflects similar docking performance to other clusters.

HADDOCK docking of CD9 with Gabrb3 yielded a top-ranked cluster (Cluster 63_1) with a HADDOCK score of –101.6 ± 13.3, indicating a moderately strong predicted interaction. The cluster size was 2, with a low RMSD of 20.9 ± 0.1 Å, reflecting structural consistency among models within the cluster. Interface analysis identified several residues within 3 Å of the predicted binding surface, including Gabrb3: 74, 79, 81, 226, 295 and CD9: 8, 22, 37, 42, 78, 83, suggesting contacts likely involve extracellular loops of CD9 and the ligand-binding domain of Gabrb3 (Fig. 4G). The interaction was energetically supported by favorable electrostatic energy (–100.6 ± 24.6 kcal/mol), van der Waals contributions (–43.8 ± 5.2 kcal/mol), and desolvation energy (–37.6 ± 3.3 kcal/mol). The buried surface area was 1393.7 ± 137.6 Å², consistent with a stable protein–protein interface. No restraint violations were detected, and the Z-score of 0.1 suggests that this docking solution is comparable in quality to other clusters within the run.

HADDOCK docking of CD9 and GAD1 identified a top-ranking cluster (Cluster 92_1) with a HADDOCK score of –120.0 ± 4.6, indicating a stable interaction. The cluster had a small size (n = 2) and an RMSD of 34.3 ± 0.2 Å, suggesting variability within the cluster but consistent scoring. Key interacting residues within 3 Å included GAD1: 266, 274, 329, 333, 397, 421, 438, 443, 445 and CD9: 5, 6, 8, 17, 20, 148, 192, 201, 205, 223, indicating both extracellular and cytoplasmic involvement (Fig. 4J). Energetically, the interaction was characterized by strong electrostatic contributions (–310.2 ± 93.9 kcal/mol), moderate van der Waals energy (–45.3 ± 6.6 kcal/mol), and buried surface area of 2124.0 ± 68.3 Å², consistent with a plausible protein– protein interface. No restraint violations were observed. The Z-score of 0.1 suggests the cluster performed similarly to others in the docking run, albeit with favorable interaction parameters.

## Discussion

### Proteomic interactions of neuronal EVs highlight their role in key synaptic processes

Our characterization of the neuronal EV interactome revealed that neuronal EV mediated proteomic interactions are tightly coupled to synaptic function and neuronal excitability machinery. In the wild-type EV interactome, we observed the enriched for canonical synaptic proteins involved in both exocytic and endocytic pathways, highlighting a close mechanistic link between EV biology and synaptic vesicle cycling. Notably, trafficking proteins such as Septin11 were enriched and are known to play multifaceted roles in synaptic and vesicle dynamics across both excitatory and inhibitory neurons ^34^. Importantly, Septins have also been shown to modulate vesicle trafficking, particularly in neurons, where they help coordinate the transport and docking of synaptic vesicles at the presynaptic terminal ^35^. Supporting this functional overlap, Snap25, a core SNARE complex protein essential for synaptic vesicle fusion and neurotransmitter release ^36^, was also detected, suggesting that neurons may repurpose synaptic fusion machinery to mediate EV release. Additionally, Bin1, a BAR domain protein that promotes membrane curvature and scission during clathrin-mediated endocytosis ^37^, emerges as a plausible effector of EV budding. Although these proteins are classically associated with neurotransmission, their selective enrichment in neuronal EV interactome suggests that neuronal EVs may exploit canonical synaptic vesicle cycling machinery for biogenesis and secretion.

We next investigated potential mechanisms underlying EV uptake by recipient neurons. Among the top-ranked enriched proteins across both healthy and amyloid brain microenvironments was Ncam1, a neuronal cell adhesion molecule well known for its roles in synaptic stabilization, plasticity, and circuit remodeling ^38–40^. Ncam1 was uniquely enriched in the biotin-supplemented condition (TB), suggesting its labeling occurs in extracellular or postsynaptic regions where exogenous biotin is required for proximity labeling. Given that target neuronal EV endocytosis remains poorly understood, we sought to validate whether Ncam1 is associated with neuronal EVs. Proximity ligation assays of cortical regions revealed significant biotinylation of Ncam1 around transfected neurons, suggesting its incorporation into neuronal EVs. In addition, *in silico* docking simulations demonstrated a stable interaction interface between Ncam1 and CD9, supporting the idea that Ncam1 may directly associate with neuronal EV membranes. Given its adhesive and signaling functions at synapses, Ncam1 may enable EVs to engage with recipient neuronal surfaces, particularly at synaptic or perisynaptic sites, serving as a molecular bridge to facilitate targeted EV docking and uptake.

While synaptic proteins involved in vesicle trafficking were expectedly among the most enriched, many of the top-enriched proteins in both healthy and AD pathological conditions were components central to neuronal excitability--including ionotropic receptor complexes, and ion transporters--revealing an interesting convergence between neuronal EV cargo and molecular machinery governing neuronal signaling and excitability. Notably, GABAergic signaling consistently emerged as one of the most enriched pathways in our analysis with nearly ten distinct GABA_A_ receptor subunits enriched, including Gabrb3, Gabbr2, and Gabra1, along with GABA recycling machinery, Slc6a1(GAT1), across both healthy and amyloid pathological conditions. These findings are particularly compelling considering previous reports that excitatory ionotropic receptor subunits, such as Gria2, can be incorporated into EVs derived from cultured cortical neurons ^41^. In contrast, only a limited number of inhibitory GABA_A_ receptor subunits have been previously detected in EVs (Gabrb3 and Gabrb2) and these were identified in non-neuronal contexts, such as human urine ^42^or hepatocellular carcinoma cells ^43^, respectively. To our knowledge, this study provides the first *in vivo* evidence that neuronal EVs preferentially incorporate a broad range of GABA_A_ receptor subunits, including several not previously reported in any EV dataset ^44^. This could suggest a previously unrecognized role for neuronal EVs in modulating inhibitory synaptic transmission through the selective packaging of ionotropic receptor subunits, potentially directing them toward lysosomal degradation pathways or enabling their delivery to target cells.

Additionally, the significant enrichment of chloride-ion transporters such as Slc12a6 suggests that EVs may play a more active and complex role than simply shuttling ionotropic receptor subunits. Beyond their well-established role in regulating neuronal chloride gradients, chloride transporters are increasingly recognized as key players in vesicular trafficking. Chloride flux is essential for maintaining the electrochemical gradients that support vesicle acidification, membrane potential regulation, and efficient cargo loading and sorting within intracellular compartments ^45^. Moreover, recent work has shown that EVs can harbor functional voltage-gated ion channels and transporters, which act as gatekeepers of ionic homeostasis, helping vesicles adapt to extracellular ionic shifts and maintain structural and functional integrity during transit ^46^. In this context, the presence of chloride transporters in neuronal EVs raises the possibility that these vesicles may not only deliver receptor complexes but also actively establish or regulate ionic gradients within their lumen or at recipient membranes. Through these dual roles, chloride transporters could influence both the composition and functionality of EV cargo and the electrophysiological properties of recipient neurons, offering a multifaceted mechanism for modulating vesicle trafficking, and membrane excitability.

### Neuronal EV interactome in amyloid pathology and membrane excitability machinery

We next examined the neuronal EV interactome in APP^NLGF^ mice to determine how amyloid pathology affects neuronal EV processes. Similar to the WT interactome, APP^NLGF^ EVs were enriched for proteins involved in vesicle trafficking, synaptic function, and neuronal excitability. Notably, several proteins with known roles in AD were enriched, including Mapt (Tau), Snca (α-synuclein) and Sncb (β-synuclein), all of which have been shown to associate with EVs in AD and related disorders, where they may contribute to the propagation of pathological protein aggregates across neuronal networks ^47,48^. Their presence neuronal EVs *in vivo* further supports the hypothesis that neuronal EVs serve as vehicles for the spread of misfolded proteins in disease contexts. Alongside these disease-linked proteins, we identified synaptic components including Ncam1, Snap25, and Vamp2, underscoring the deep functional overlap between EV secretion, uptake and the synaptic vesicle machinery. This mirrors our observations in the WT interactome, where canonical elements of neurotransmitter release also appear to support the biogenesis, release and uptake of neuronal EVs.

We also observed robust enrichment of inhibitory signaling components in the APP^NLGF^ EV interactome, including multiple subunits of the GABA_A_ receptor complex, Gabrb3 and Gabra2, as well as the GABA transporter Slc6a1 (GAT1). While similar enrichment of GABAergic signaling proteins was observed in the WT interactome, supporting a conserved role for EVs in inhibitory synaptic function, this pathway takes on particular significance in the context of AD where GABAergic dysfunction and resulting network hyperexcitability are well-documented features of early AD ^49–51^. Recent studies have shown that brain-derived EVs in AD preferentially accumulate in inhibitory neurons ^48^, consistent with our observation that neuronal EVs interact with inhibitory neuronal markers in both healthy and AD pathological conditions. Notably, Gad1, identified in our shared enriched protein STRING network, is a GABA-synthesizing enzyme uniquely expressed in inhibitory neurons. It showed significant enrichment in the APP^NLGF^ brain compared to saline controls, but not in the WT condition. Because the fold change was modest, we validated that neuronal EVs interact differentially with Gad1 using proximity ligation assays (PLA). Supporting this, HADDOCK-based protein docking revealed a potential direct binding interface between CD9 and Gad1. These findings suggest a previously unrecognized mechanism by which EVs may contribute to GABAergic dysfunction in AD, potentially through aberrant interactions that lead to the mislocalization or lysosomal degradation of essential proteins such as Gad1. This mechanism could disrupt GABA synthesis and contribute to the early network instability and excitability changes observed in AD. Further investigation is warranted to determine whether targeting EV–GABAergic molecular interactions may offer therapeutic potential for restoring inhibitory balance in AD.

Lastly, while both datasets showed enrichment of voltage-gated potassium (K⁺) channels, the APP^NLGF^ EV interactome was particularly distinguished by a broader diversity of uniquely enriched K⁺ channels and transporters. Core regulators of membrane excitability, including Kcnma1 (BK channel) and Kcna2, as well as modulatory proteins such as Dpp6, were consistently enriched across both healthy and AD conditions. While previous studies have demonstrated that EVs can carry voltage-gated ion channels, recent findings show that these channels may remain functionally active within the EV membrane, contributing to ionic homeostasis, osmotic regulation, and potentially signaling capacity ^46^. Kcnma1, one of the most highly enriched proteins in our dataset, has been shown to support volume regulation and membrane potential adaptation in EVs during transit. The distinct composition of K⁺ channels and transporters in APP^NLGF^ EVs, including proteins not previously reported in EVs such as Kcnk2, may reflect a selective remodeling of EV cargo in response to amyloid-associated ionic imbalance. This shift could represent an adaptive mechanism, buffering osmotic stress during EV biogenesis or enabling targeted modulation of excitability in recipient neurons. The observed increase in both K⁺ channels, transporters and modulators may therefore act as a compensatory response to hyperexcitability and disrupted ion homeostasis in the AD brain. However, functional validation is needed to resolve whether these changes in EV K⁺ channel content actively contribute to the modulation of neuronal excitability or instead represent a passive consequence of membrane remodeling in a diseased microenvironment.

## Acknowledgments

We thank all current and former members of the Laboratory of Molecular NeuroTherapeutics for their technical assistance. This work was supported by the NIH (RF1AG054199 to T.I., R01AG054672 to T.I., R01AG066429 to T.I., R01AG067763 to T.I., R01AG072719 to T.I., RF1AG082704 to T.I. and S.I., R01AG079859 to S.I.), and Cure Alzheimer’s Fund (to T.I. and S.I.).

## Authors’ contributions

T.I. Conceptualized the study; T.I. and S.I. supervised the study. Y.C., M.A., and B.G. developed the experimental design, developed methodologies, and performed experiments. B.G. and Z.L. performed data and bioinformatic analysis, data interpretation, and prepared figures; Y.C. and A.L.P. performed stereotaxic surgeries, biotin injections; J.M.K. performed AlphaFold, HADDOCK analysis, biotin injection and brain sectioning; B.M. and Y.C. performed brain sectioning, immunohistochemistry, microscopic imaging, and quantifications; S.N. and M.P.G.P. performed mass spectrometry and data analysis; B.G., Y.C., S.I., and T.I. wrote and edited the manuscript. All the authors have read, edited, and approved the final version of the manuscript.

## Ethics declarations

The authors declare no competing interests.

