## Supplementary figures for "Proteomic characterization of neuronal extracellular vesicle interactomes in Alzheimer’s disease mouse model through TurboID-based proximity labeling"

### Supplementary Information

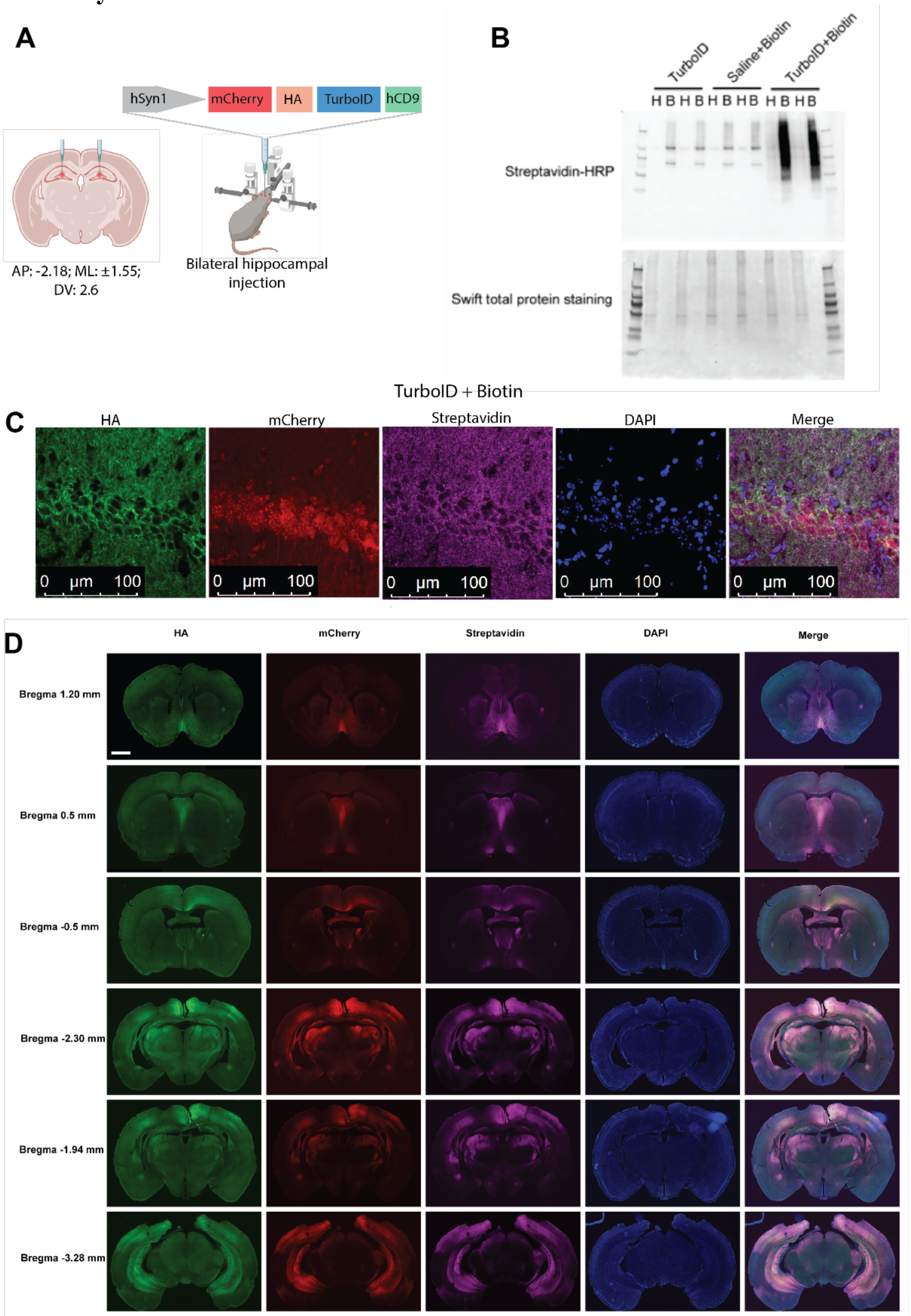


**Supplementary Figure 1: Generation and Validation of TurboID-CD9 Injection into the Mouse Brain**

**(A)** Intracranial injection of AAV TurboID vector. **(B)** Western blot analysis confirming the biotinylation of proteins bound to streptavidin beads in samples from TurboID+Biotin group (H:

homogenate, B: biotinylation fraction sample). **(C)** Confocal microscopy of immunofluorescence staining of HA, mCherry and Streptavidin in the TurboID+Biotin group. **(D)** Whole-brain immunofluorescence staining of HA, mCherry and Streptavidin in different regions of the TurboID+Biotin group.

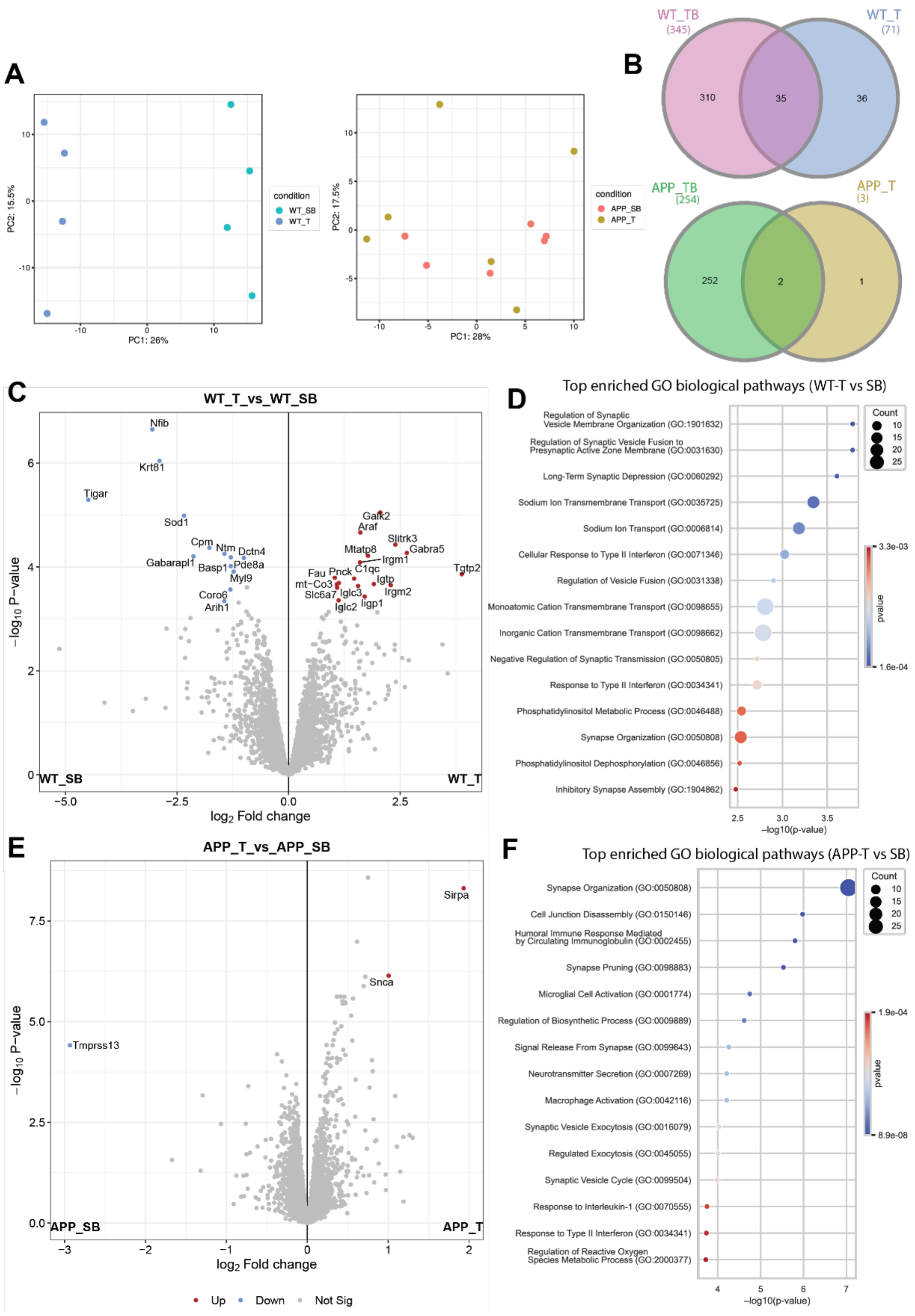


**Supplementary Figure 2: Comprehensive proteomic analysis of experimental control groups (A)** Principal component analysis (PCA) of the T group against the SB group in WT and APP mice. **(B)** Overlap of enriched proteins between WT and APP TB groups (TBvsSB) and T controls (TvsSB). **(C)** Volcano plot showing enriched proteins in the WT T group compared to SB control. Comparison includes WT-T vs. WT-SB, with significance thresholds set at one-way

ANOVA, FDR < 0.05, log2 FC > 1, and log2 FC < -1. (**D**) GO biological pathway analysis of enriched proteins in WT T vs SB groups. **(E)** Volcano plot showing enriched proteins in the APP T group compared to SB control. Comparison includes APP-T vs. APP-SB, with significance thresholds set at one-way ANOVA, FDR < 0.05, log2 FC > 1, and log2 FC < -1. (**F**) GO biological pathway analysis of enriched proteins in APP T vs SB groups.

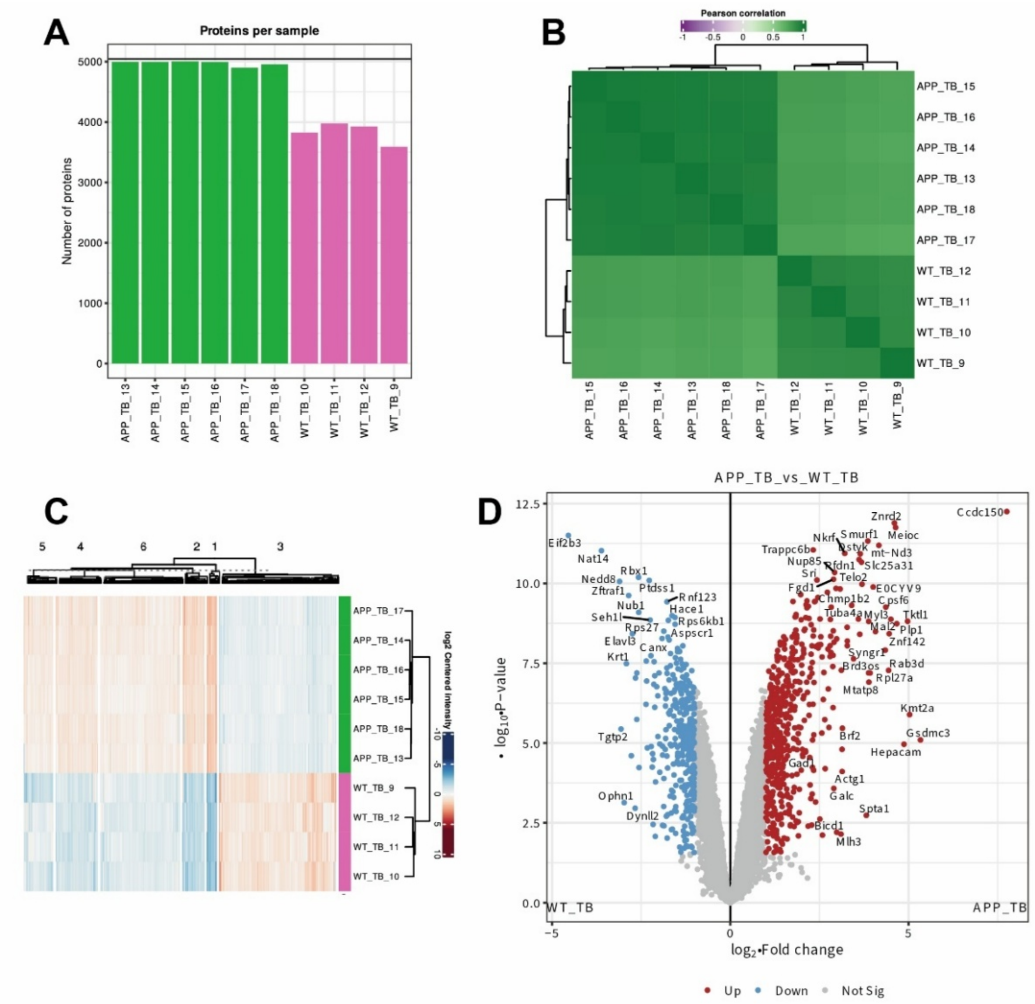


**Supplementary Figure 3: Proteomic comparison of CD9 Interactome in APP$^{NLGF}$ Mice vs. WT Mice (A)** Number of proteins detected in each of our TB groups. **(B)** Pearson correlation plot of our TB groups. **(C)** Heat map of protein expression across the TB groups. (**D**) Volcano plot showing enriched proteins in the APP-TB group compared to WT-TB. Comparison includes APP-TB vs. WT-TB, with significance thresholds set at one-way ANOVA, FDR < 0.05, log2 FC > 1, and log2 FC < -1.
